# No overlap in neurophysiological predictors of moment-to-moment fluctuations of behavioural variability and subjective attentional state

**DOI:** 10.64898/2026.07.30.741808

**Authors:** Marlou Nadine Perquin, Marcus Daghlian, Gavin Perry, Krish D Singh, Aline Bompas

## Abstract

Behaviour fluctuates from moment to moment – often attributed to changes in subjective attentional states (e.g., mind wandering). Yet, it remains unclear how closely behavioural variability and subjective attentional state are related, and whether they share biological mechanisms. Using ∼2 hours of Metronome Response Task data, we combined magnetoencephalography, eye and head tracking, pupillometry, and thought probes to predict trial-to-trial fluctuations in both. Although they covaried weakly, this relationship was largely accounted for by time-on-task. Several markers including blink duration, pupil variability, and head movement predicted both measures. Dissociations also emerged: behavioural variability was linked to broadband functional connectivity and delta power, whereas off-taskness was predicted by theta, alpha, and beta power. Critically, predictive strength of behavioural variability and subjective reports were uncorrelated at the individual level, with substantial individual differences. For both outcomes, multimodal models outperformed univariate analyses. Our findings show that behavioural variability and subjective attentional state are largely dissociable, challenging the assumption that they share underlying process or that one causes the other.

## Introduction

Variability over time is ubiquitous in any repetitive human behaviour: even under controlled laboratory conditions, and no matter how straightforward the task, reaction time (RT) varies from trial to trial and errors occur. Only a fraction of this RT variability can be explained by external factors, such as task conditions or time-on-task (i.e., trial number or elapsed time within the session) or by recent behaviour (Perquin et al., 2023; 2024). The rest is assumed to 1) originate from spontaneous fluctuations in brain-body systems (Faisal et al., 2008; Urai, 2025), and 2) manifest subjectively through fluctuations in metacognitive states (e.g., Seli et al., 2013; Smallwood & Schooler, 2015). The present work explores these assumptions and their interplay.

To the extent that “spontaneous” is taken to mean “without identifiable external causes”, assumption 1 is hardly debatable, but the plethora of potential sources of physiological within-subject variability, overlapping at different spatial and temporal scales, presents a large methodological challenge – especially for non-invasive human research. Numerous neurological and physiological markers have been associated to task performance, including electroencephalography (EEG) features (Gonzales-Andino, 2005), functional magnetic resonance imaging (fMRI) features such as BOLD response in task-related brain areas and the default mode network (Kucyi, 2016), or pupil diameter (Unsworth & Robinson, 2016). Yet methods vary considerably across studies even within single modalities, both in tasks employed and in features examined. Technical and financial limitations constrain researchers to focus on one (or a few) biological measures at a time, each only explaining a small part of behavioural variance (Duffy et al., 2024). It therefore remains essentially unknown to what extent their predictive powers overlap or complement each other.

Assumption 2 intuitively relies on the idea that cognitive resources intermittently shift away from the task, leading to bouts of poor performance and reduced subjective on-taskness (i.e., the degree to which a person feels focused on the task rather than engaged in task-unrelated thoughts) or “mind wandering”. This assumption is supported by empirical observations of within-subject correlations between objective performance and subjective on-taskness across time or trials during tasks involving tapping (Seli et al., 2013, Laflamme et al., 2018; Perquin et al., 2023), sensory detection (MacDonald et al., 2011; Jin et al., 2019), sustained attention to response (SART; Qin, Perdoni & He, 2011; Jin et al., 2019; Robison & Unsworth, 2016), continuous performance (CPT; Bertschi et al., 2026), reading comprehension (Schooler, Reichle & Halpern, 2004; Ward & Wegner, 2013), and driving (Baldwin et al., 2017; Pepin et al., 2020). Crucially though, these correlations are weak, explaining only between 2-7% of the variance (Perquin et al., 2023; Kane et al., 2021) – insufficient to support the idea that poor performance and subjective off-taskness are two sides of the same coin indexing objective and subjective mind wandering (Martinez-Perez et al. 2023), but consistent with a shared susceptibility to factors such as time-on-task (Zanesco et al., 2024).

Biological correlates of objective performance and subjective on-taskness show some overlap (Andrillon et al., 2021; Zanesco et al., 2021) alongside clear dissociations (Whitmarch 2017; Qin 2011; Kucyi 2016; Zuberer et al. 2021). RT variability likely results from many sources of internal noise, an intrinsic and necessary feature present throughout the entire sensory and nervous system (see Ermentrout et al., 2008; Faisal et al., 2008; Garret et al., 2013; Renart & Machens, 2014; Waschke et al., 2021 for comprehensive reviews), many of which may not be consciously accessible. Reciprocally, metacognitive judgments may also arise from multiple sources, only some of which may directly interfere with behaviour. If attentional state contributes only a small proportion of overall RT variability (the proportion accessible for conscious reporting), then attentional state and RT variability may only be loosely related biologically. Too few studies investigated the neural bases of contingent objective and subjective measures of performance to estimate their (dis)similarities, and none have conducted within-subject analyses of overlap.

Rather than being treated as noise, behavioural variability is increasingly being recognised as conceptually informative. It intersects with key questions about determinism and exploration–exploitation dynamics (Carpenter, 1999; Duffy et al., 2024; Perquin et al., 2024; Uddin, 2020; Urai, 2025). Behavioural variability is a reliable individual trait across different contexts and modalities (Hedge et al., 2018; Hultsch et al., 2000, 2002; Perquin & Bompas, 2019; Perquin et al., 2023; Saville et al., 2011, 2012). Its potential link with metacognition relates to awareness of one’s own biological states. At the applied level, physiological measures and subjective sampling of off-taskness have both been proposed as solutions for detecting or predicting, and therefore reducing, episodes of poor performance. This reduction appears particularly desirable in high-risk contexts such as driving and air traffic control. This itself relies on a 3^rd^ assumption: that human beings are typically poor at assessing their current performance or their risk of errors (Schmidt, 2009). Assumption 3 appears to originate mostly from early vigilance studies manipulating factors such as fatigue and sleep deprivation, and reports that drivers do not seem to realise when they are at risk of falling asleep or crashing (Belz et al., 2004; Moller et al., 2006; Philip et al., 1997; 2003). While experience sampling literature has vastly increased over the past 15 years, only few studies ask participants how well they think they performed (Whitmarch 2017; MacDonald 2011).

In the current study, we address these key assumptions through a multi-modal approach combining magnetoencephalography (MEG), eye and head tracking, pupillometry and subjective reports of off-taskness and perceived performance. We also consider short-term and long-term temporal trends. Despite a rapidly growing corpus of published studies co-investigating objective and metacognitive states (> 4000 in Dimension^1^), or the biological markers of either of these, none have comprehensively investigated all these ingredients. Furthermore, while most literature relies on EEG or fMRI, their results will be limited by poor signal to noise ratio and poor spatial resolution for the former, and poor temporal resolution for the latter. In contrast, MEG offers quantification of dynamic neural activity with millisecond temporal resolution functional connectivity across cortical networks (Baillet, 2017) – allowing for examination of moment-to-moment fluctuations in neural activity. In this study, we estimated the trial-to-trial fluctuations in oscillatory power and functional connectivity across 90 brain regions.

We used the metronome response task (MRT; Seli et al., 2013), which offers a single, semi-continuous, outcome measure of behavioural consistency. Our participants heard a tone every three seconds and were instructed to press a button in synchrony with the tone, while occasional pseudo-random thought probes asked them to rate their off-taskness and perceived performance just prior to the probe (Weinstein, 2018). This task presents two key advantages: 1) it does not rely on alternating across empirical conditions (e.g. Go and Nogo stimuli), therefore avoiding external variance that would confound within-participant variance; 2) there are no errors, and it provides no implicit nor explicit feedback on performance, so these cannot influence upcoming behaviour or subjective reports. The objective measure of performance is the standard deviation of “reaction times” over 5 taps (i.e., over a 15 seconds window, thereafter SDRT). Our aim is to provide the first comprehensive capture of brain and body markers of within-participant fluctuations in objective performance and subjective attention, and their overlap. While previous studies have relied mainly on limited within-participant repetition with larger participant number, we adopt the opposite approach, opting for deep within-subject sampling: our final sample consists in 21 participants, but the core within-participant analyses rely on 1440 trials (∼2 hours of task), which is essential to capture within-participant fluctuations. We predict within-subject fluctuations in behavioural variability and off-taskness from three predictor groups: MEG (delta, theta, alpha, beta, low-gamma and high-gamma power, and functional connectivity), overt physiology (mean pupil size, pupil variability, blink rate, blink duration, and head movement), and task-derived measures (subjective performance ratings, time-on-task, and previous behaviour).

## Results

### Many measures show weak associations with SDRT and off-taskness ratings

First, we correlated SDRT over the 15 seconds before each probe with subjective off-taskness ratings. One Pearson r-value was obtained for each participant, and the group distribution of r-values was compared to zero. Within-subject correlations were overall positive, indicating that participants were more variable during more off-task states (Figure 1, black violin). Consistent with Perquin et al. (2023), the group result was strong (BF_10_ = 23) yet relied on very small effect sizes: the group median of shared variance was only 2.0%.

**Figure 1.**
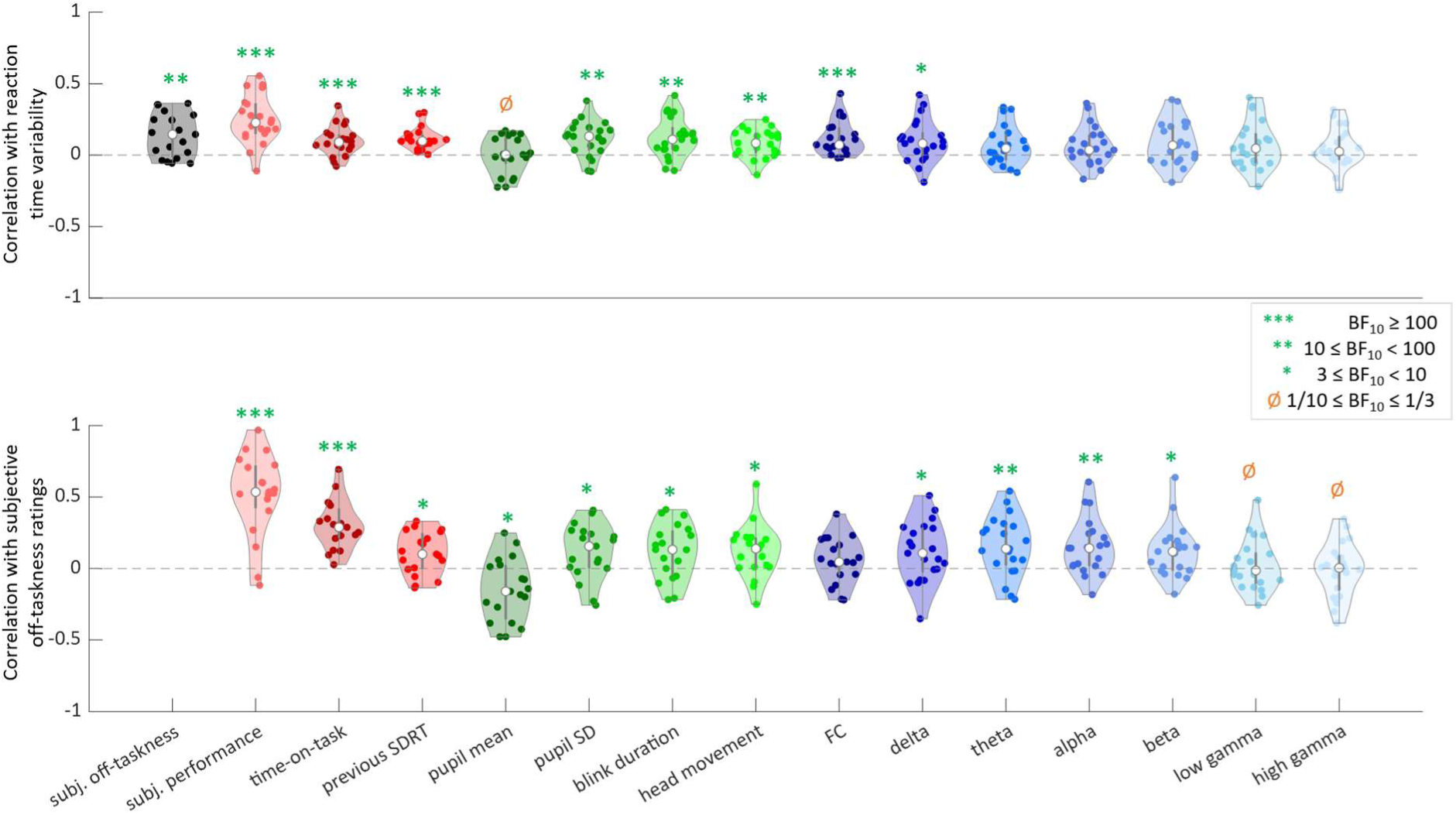
Violin plots of correlation coefficients (Pearson’s r-values). Each coloured point shows the r-values of one participant, with the median across the group in white. The black violin shows the correlations between reaction time variability and subjective off-taskness. Each other violin reflects the correlations between each variable and 1) reaction time variability (top panel) and 2) off-taskness (bottom panel) window. Task-related measures are shown in red, overt physiological (eye and body) measures are shown in green, and neural measures are shown in blue. Evidence for the alternative hypothesis (i.e., the distribution is different from zero) is indicated by green stars (BF10 > 3), evidence for the null-hypothesis by orange null signs (BF01 > 3), and indeterminate evidence by the lack of a symbol. Note that subjective performance is reverse-coded such that low values indicate better performance, hence the positive correlation with subjective off-taskness and SDRT (see methods).

Using a similar approach, we assessed the independent relationship of each predictor with SDRT and off-taskness (Figure 1). Many distributions were clearly statistically above zero. Overall, the numerically strongest predictors were task-related (red violins on Figure 1) rather than biological measures (green and blue violins). Subjective performance ratings showed the highest correlation with both SDRT and off-taskness (5.3% and 30% explained variance respectively). Time-on-task and SDRT on the previous window correlated relatively well too with both current SDRT and off-taskness.

Among the MEG measures (blue violins), SDRT only correlated with functional connectivity and delta power (1-4 Hz), while off-taskness did not correlate with functional connectivity but did correlate with oscillatory power throughout the 1-30 Hz range. Three overt physiological measures (cumulative blink duration, standard deviation of pupil diameter and head movement, green violins) numerically outperformed the best MEG measures. Overall, many r-values were very low but with high Bayes Factors, providing confidence that variables are positively, albeit weakly, correlated. Considerable inter-individual variability was evident, with no predictor consistently influencing all participants; all violins overlapped with, or came very near, the zero line.

#### Whole-brain versus network-specific activity

Above, we report whole-brain connectivity. We also ran network-specific analysis, focusing on nodes corresponding to key components of the DMN, and 2) task-related auditory and motor nodes. Both network-specific analyses showed the same directional effect as the whole-brain analysis, but with numerically lower shared variance. To determine if the increase in connectity was associated with a limited set of connections, we correlated the strength of each connection separately with both variability and off-taskness (2*4005 correlations in total). At the group level, most correlations were positive and many reached significance (RT variability: 93.6% positive, 42.7% significant; attentional state ratings: 74.9% positive, 15.1% significant). The distributions were approximately symmetric and showed no clustering within canonical networks, suggesting that the observed effects reflect a widespread increase in connectivity. Importantly, there was little overlap between the connections associated with RT variability and those associated with attentional state ratings (see Supplementary Materials). Similarly, region-specific analyses of oscillatory power did not identify any particular networks or regions as key drivers of the effects, and again revealed minimal overlap between the two outcome measures.

### Predictors of SDRT and off-taskness do not overlap in most people

Above, we examined objective and subjective outcomes separately. However, if they are both manifestations of the same mechanisms, their predictors would be expected to largely overlap at the individual level, even if these predictors vary largely across individuals. To quantify this overlap, we plotted the absolute correlation coefficient of each predictor with SDRT against its absolute correlation coefficient with off-taskness (Figure 2).

**Figure 2.**
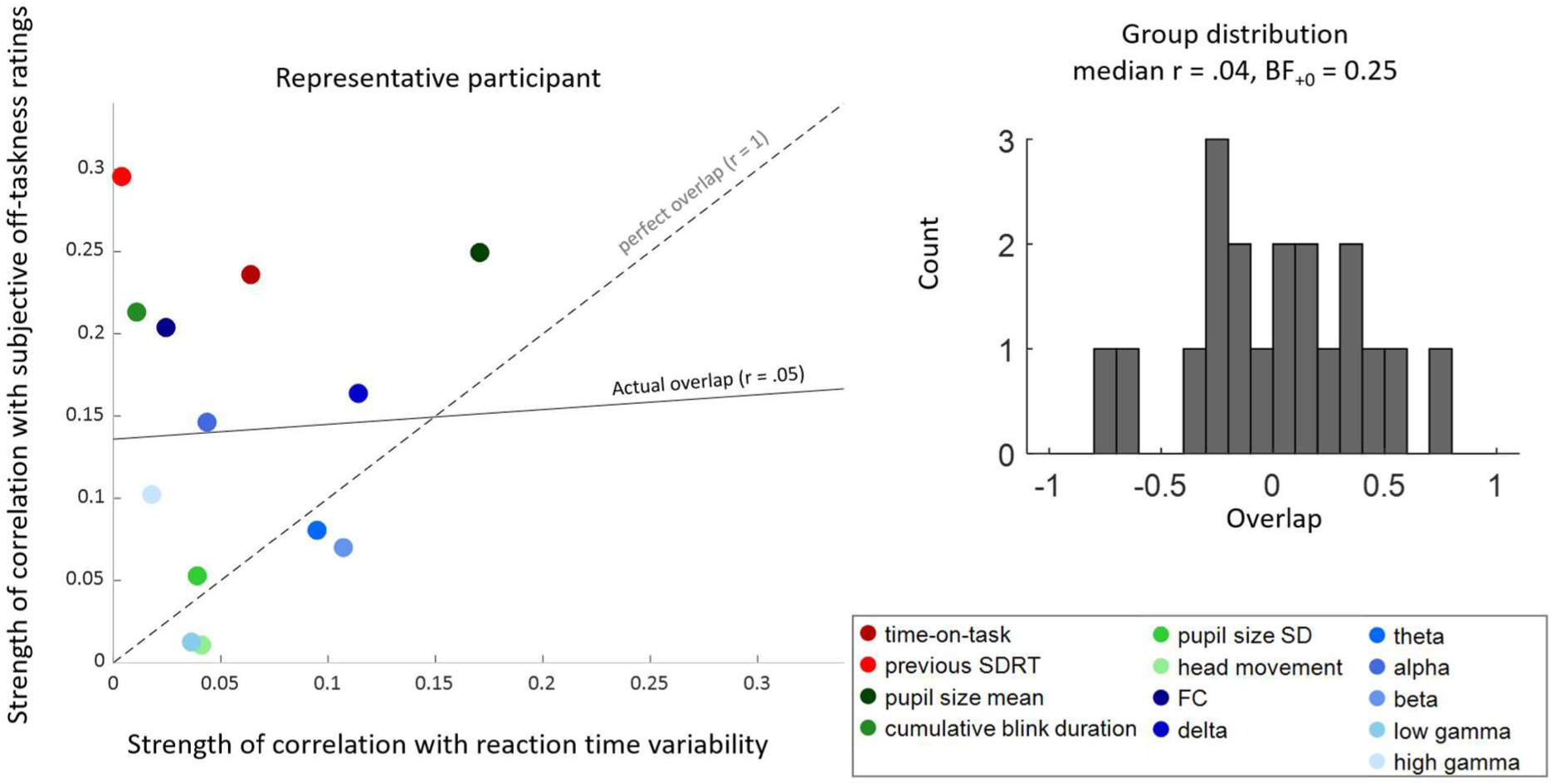
No overlap between the predictors of RT variability and off-taskness. **Left**. The single-participant example shows the absolute r-values between each predictor and 1) RT variability (x-axis), and 2) off-taskness (y-axis). Predictors include all those from Figure 1, except subjective performance (randomly collected just before or just after off-taskness, hence not a “predictor”). The dashed line reflects a perfect overlap between the predictors (correlation of r = 1) while the full black line shows the actual linear correlation (r = .05 in this example participant). **Right.** The distribution of the linear overlap across participants, which did not differ from zero.

A strong overlap would be reflected in a similar ordering of predictors along both axes. However, for the participant shown in Figure 2, the relationship between the two sets of correlations was near zero (*r* = .05; solid line), indicating little to no overlap between the predictors of SDRT and off-taskness (see Supplementary Materials for all participants seperately). At the group level, the distribution of r-values was not higher than zero (BF_+0_ = 0.25), with the median r-value on the group level being near zero (0.04). Accordingly, the participant presented in Figure 2 exhibits a slope that is closest to the group median.

### Many measures correlate with many measures

So far, we have been analysing the predictors in isolation, but it is unlikely that they are independent from each other. Indeed, we found that many predictors showed small correlations with many other predictors (Figure 3), most of which positive. Again, we found conclusive but weak relationships: the Bayes Factors are high, but the effect sizes are very low. Unsurprisingly, the correlations were strongest among similar measures: task-measures to other task-measures (red box on Figure 3), overt physiology to overt physiology (green box), and MEG measures to MEG measures (blue box).

**Figure 3.**
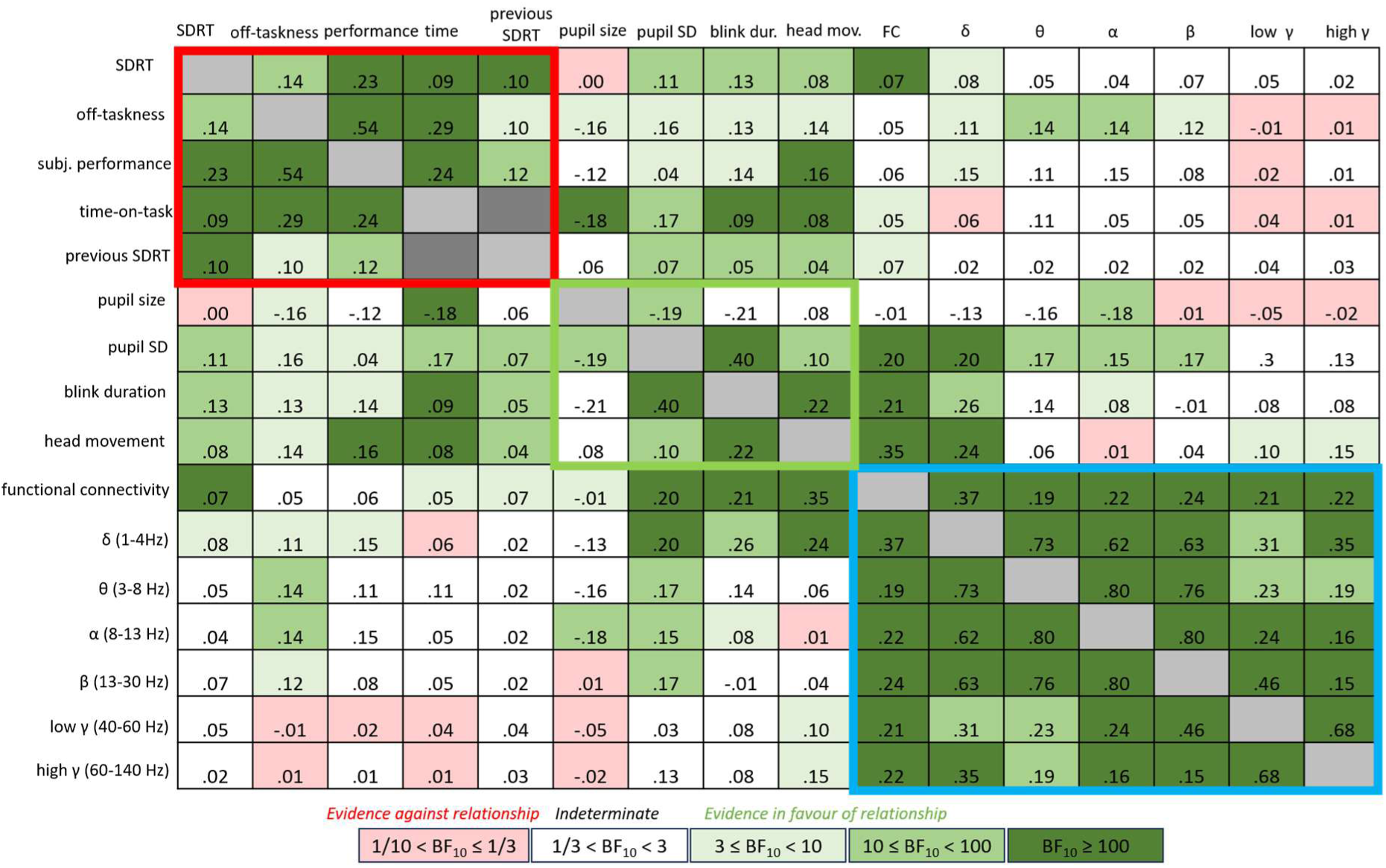
Correlation matrix across all measures. Numbers in the table reflect the group median of the Pearson r correlation coefficients, providing some indication of the strength of the correlation. Colours of the boxes indicate the Bayes Factors resulting from the two-sided one-sample t-tests, testing if the group distribution is different from zero. Evidence for the hypothesis that distribution is different from zero is shown in green (BF10 > 3), evidence for the null-hypothesis in red (BF01 > 3), and indeterminate evidence in white. The coloured boxes (red, green, and blue) indicate the pairs of the same types of measures (task-related, over physiological, and MEG respectively). Note that the correlation between previous SDRT and time-on-task was not computed, on the assumption that previous behaviour cannot cause time to pass by.

We find strong correlations across the MEG frequency bands, but comparatively limited and inconsistent relationships with neurophysiological markers – except for FC and delta which correlated with head movement and eye measures. In fact, the correlation between functional connectivity and head movement (r = 0.35) was the strongest correlation in the table outside of the coloured boxes.

### Controlling for time-on-task reduces predictor-outcome relationships

Many of the variables were affected by time-on-task, which can impact predictor-outcome relationships. In particular, time-on-task had a systematic detrimental influence on subjective off-taskness and perceived performance, more so than on objective performance. To control for these temporal effects, we tested if predictor-outcome relationships would change after detrending all variables. We found that all but one of the 27 distributions shifted closer to zero (Figure 4). Notably, the correlation between SDRT and off-taskness reduced substantially. While the distribution was still statistically above zero, the median shared variance dropped from 2% to < 1%.

**Figure 4.**
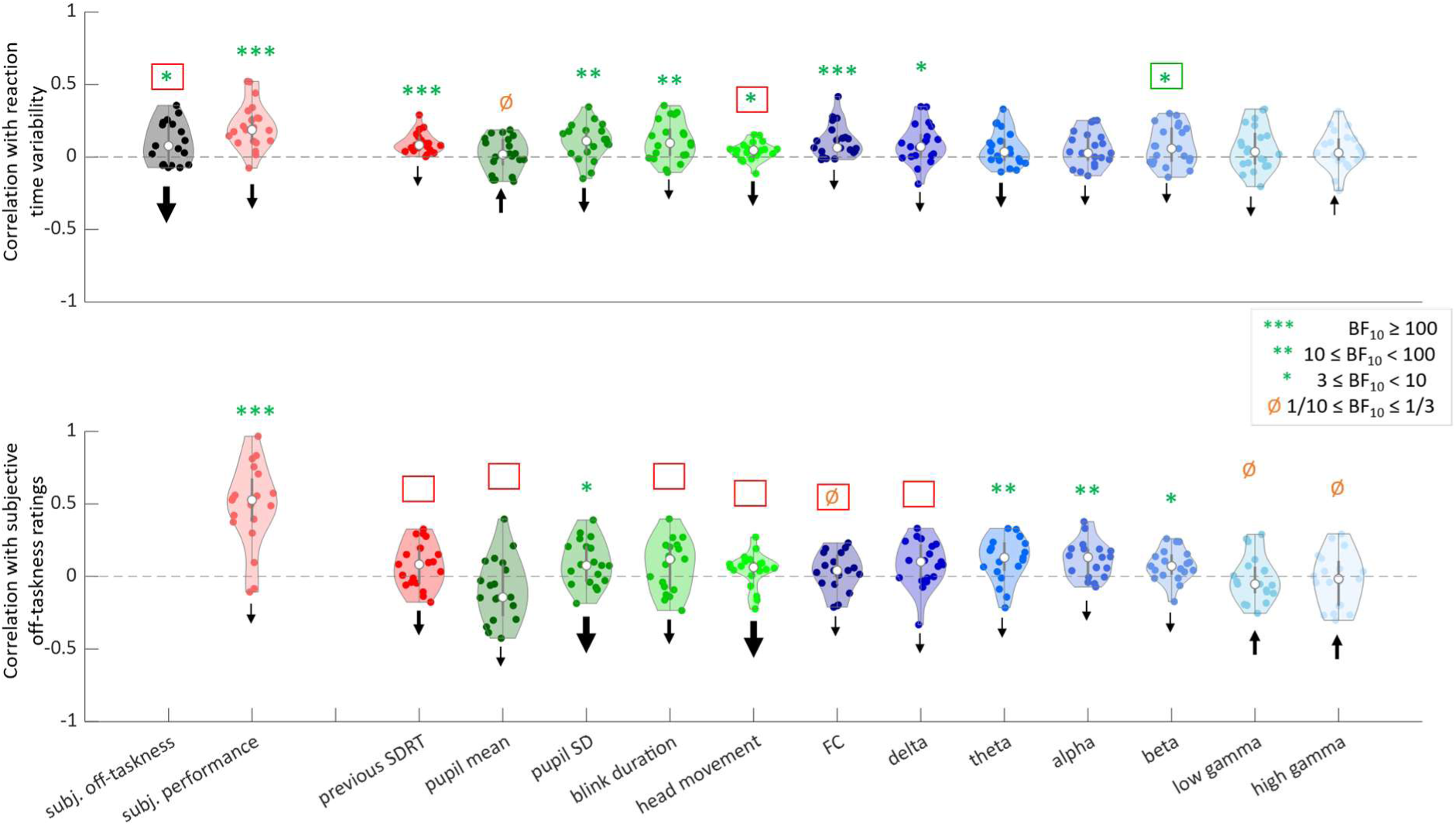
Violin plots of correlation coefficients (Pearson’s r-values) after detrending both predictor and outcome variables. Conventions are the same as in Figure 1. Black arrows indicate whether difference in the group median between original and detrended correlations is negligible (< .01, thin), medium (> .01 but < .05, medium width), or substantial (> .05, thick). The coloured boxes around the Bayesian evidence indicate that the BF_10_ has changed categories (red: BF_10_ decreased, green: BF_10_ increased).

For SDRT, detrending revealed a new relationship with beta power, but only moderate shifts otherwise that led to no changes to our overall conclusions. In contrast, several predictors of off-taskness were no longer statistically different from zero, including most of the overt physiological predictors, previous SDRT, and delta power.

### Explained variance by all predictors

Multiple regression analyses were run to quantify how much variance can be explained by combining objective predictors. Four models were run for both outcome measures, to quantify the variance explained by 1) task-related, 2) overt physiological, 3) MEG, and 4) all predictors. The estimates of explained variance were corrected for overfitting (Perquin et al., 2024). Overall, the explained variance of off-taskness was almost three times as high as for SDRT – with all predictors combined, we could explain almost 1/3 of the total variance across individuals (Figure 5, right panels). However, there were large differences between individuals. The MEG predictors explained the most variance, though the absolute difference in explained variance is modest compared to the other two models. Because the predictor groups are correlated, they explain overlapping variance, and the explained variance of the full model is thus smaller than the sum of the variances explained by the individual predictor groups – particularly for the off-taskness ratings.

**Figure 5.**
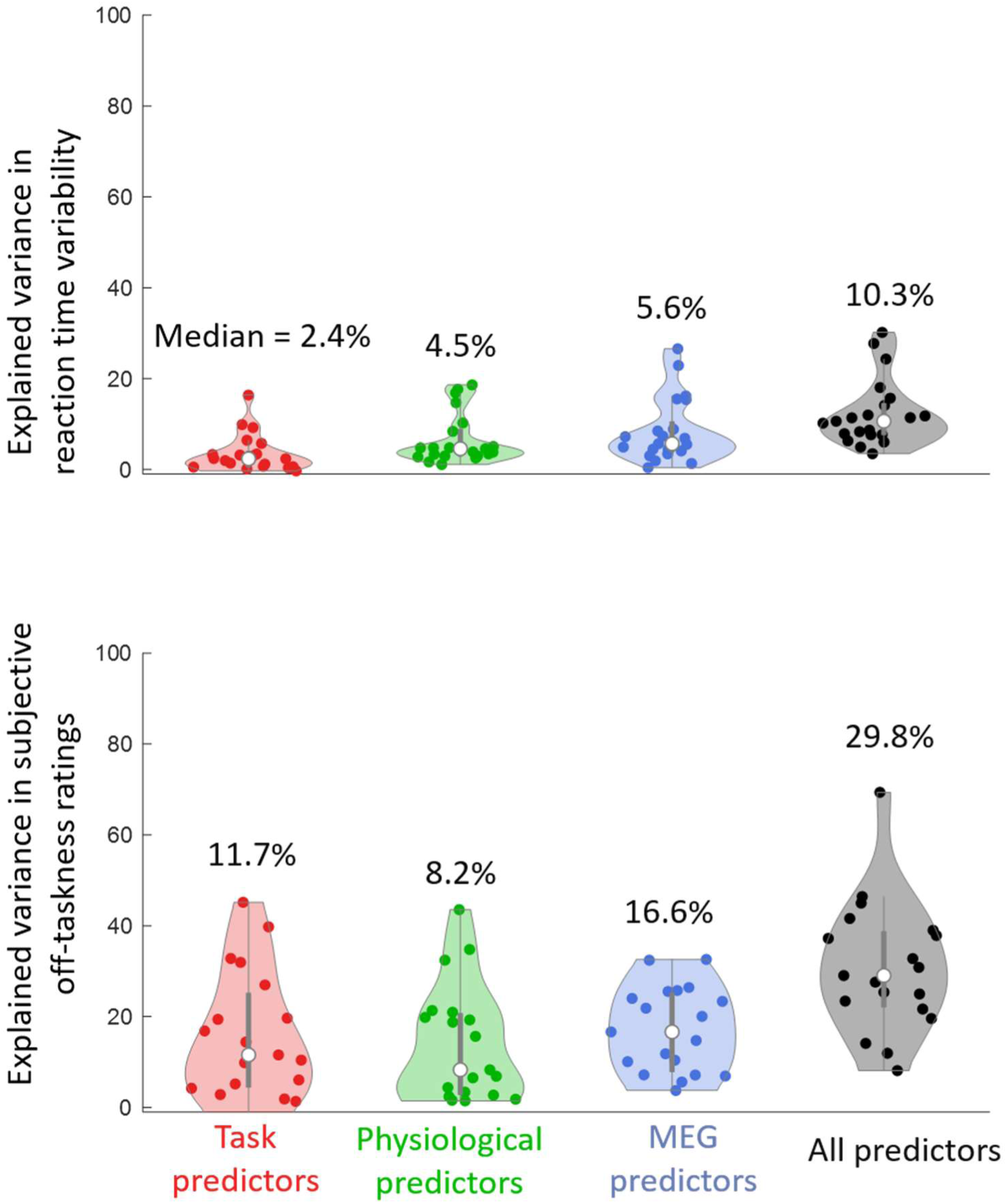
Explained variance (R^2^) in SDRT (top panel) and subjective off-taskness (bottom) by the GLMs. Each coloured dot represents one participant, with the median being represented by the white dot. For both outcome variables, four models were tested containing: 1) task-related (previous SDRT and time-on-task, and SDRT for off-taskness), 2) overt physiological (pupil size, pupil variability, blink duration, and head movement), 3) MEG (delta, theta, alpha, beta, low gamma, high gamma, and functional connectivity), and 4) all objective predictors. Note that the subjective ratings were not used as predictors. To prevent overfitting, the R^2^ values were corrected for variance explained by noise (see Methods for more details). Group medians are printed on top each distribution in text.

## Discussion

In the current study, we examined multiple behavioural, physiological, and neural predictors of behavioural variability and subjective attentional state. Our findings indicate that behavioural variability and subjective attentional state are only weakly correlated and associated with different biological markers, at the individual and group levels – echoing and strengthening previous reports questioning the assumption of a strong relationship between objective performance and subjective attentional states (Head & Helton, 2016; Perquin et al., 2019; 2021; 2023).

When considered in isolation, many predictors showed statistically convincing but uniformly weak relationships with both SDRT and off-taskness. However, predictors were themselves intercorrelated, and controlling for time-on-task attenuated most predictor–outcome relationships, particularly for attentional state. When combined, all predictors explained about 30% of variance in attentional state, but only about 10% of variance in SDRT (∼10%). These findings cast doubts on the capacity of current non-invasive biological measures to predict episodes of poor performance in tasks involving sustained attention.

### Behavioural variability and subjective off-taskness are distinct phenomena

The observed weak within-subject association between behavioural variability and subjective off-taskness aligns with prior literature (Chidharom et al., 2025; Kane et al., 2021; Perquin et al., 2020; 2023), and demonstrates that this relationship diminishes even more after controlling for time-on-task. Recent evidence from empirical manipulation further supports this dissociation: short task breaks and task switches can reduce subsequent behavioural variability without affecting subjective reports of off-task thought (Chen et al., 2025), consistent with partially separable underlying mechanisms. Between-subject dissociations have also been found: person-level estimates of behavioural variability do not correlate with attention traits or mean off-taskness (Perquin et al., 2019; 2023), and structural equation modelling approaches indicate that behavioural and subjective measures are not well captured by a single latent factor (Unsworth et al., 2021; Welhaf & Kane, 2023; 2024).

Most research has examined behaviour-attention and neural-attention relationships in isolation, leaving the three-way link underexplored (e.g., Braboszcz & Delorme, 2011; Groot et al., 2021; Jin et al., 2019; Kam et al., 2021). Studies addressing this three-way link typically find dissociable neural profiles underlying RT variability and attentional state, both in E/MEG (MacDonalds et al., 2011; Qin et al., 2011; Whitmarch et al., 2017) and fMRI (Godwin et al., 2023; Groot et al., 2022; Kucyi et al., 2016; Zuberer et al., 2021). Group-level overlap has been found for sleep-like slow waves in the delta frequency range, with distinct spatial patterns that predicted specific combinations of behavioural and subjective outcomes (Andrillon et al., 2021). Furthermore, EEG microstate analyses showed that coverage (proportion of time the signal is assigned to a microstate) and global explained variance (proportion of EEG signal variance accounted for by a microstate) of microstate C were positively associated with both RT variability and subjective attentional state (Zanesco et al., 2023). This may hint at but do not directly prove shared underlying mechanisms, as the parameters do not reflect spatial specificity or causal function, and as the microstate-outcome relationships differed substantially across individuals. Building on this literature, our current study confirms dissociations in predictors across different overt and covert neurophysiological markers while extending findings beyond alpha power. Crucially, it is the first to directly test the overlap of predictors of behavioural variability and off-taskness on the individual level. Though both shared and dissociable predictors are found on the group level, the individual-level analyses show no overlap, evidencing that underlying mechanisms are distinct.

Despite this accumulating evidence, behavioural variability and subjective attentional state are still frequently treated as two manifestations of a single attentional mechanism (e.g., Martinez-Perez et al. 2023; Qin et al., 2011). Our results rather suggest a different interpretation, at least during the metronome task: Subjective attentional state may be understood as epiphenomenal (rather than a modulating influence on behaviour), while behavioural variability arises from many sources of internal fluctuations across multiple time scales which are likely to be consciously inaccessible, unless they become extreme (e.g. waking up after falling sleep). Reported correlations between behaviour and subjective attentional states are driven by a shared susceptibility to time-on-task and the influence of performance monitoring on attention reports.

This interpretation is supported by the current findings – explaining why the shared variance between behavioural variability and subjective attentional state drops below 1% when accounting for time-on-task, and why subjective performance is a better correlate of behaviour compared to subjective attention, to the extent that behaviour and subjective no longer correlate after removing the variance of performance ratings. It is also consistent with several findings that are difficult to explain otherwise. First, Perquin et al. (2020) have shown that participants are unable to use self-pacing to reduce upcoming behavioural variability or improve performance in both psychophysical and ecological tasks, indicating that the internal states driving behavioural variability remain largely inaccessible to conscious control. Second, reports of off-taskness in the MRT increase when participants receive fake ‘incorrect’ feedback and when artificial stimulus delays are added to induce the perception of poor performance (Polychroni et al., 2025), showing that performance monitoring influences experiences of on-taskness. Third, participants’ awareness of whether a word or blank screen presented during the SART before thought probes is independent of their subjective attentional state – demonstrating that any behaviour-metacognition coupling is absent on a simultaneous task (Head & Helton, 2016).

Taken together, our findings challenge the assumption that subjective attentional state provides a reliable window onto the processes that generate behavioural variability. This distinction has important implications for applied frameworks that aim to monitor and intervene on behaviour in real-world settings such as driving or air-traffic control where neurophysiological markers of “attention” are often assumed to directly index performance-relevant internal states. Our results suggest that this assumption may be fundamentally flawed, and that interventions targeting subjective off-taskness are unlikely to yield improvements in behaviour.

### Neurophysiological predictors of behavioural variability and subjective off-taskness

#### Univariate predictors of behavioural variability

Our study fits into a large body of work examining oscillatory activity preceding behavioural performance (e.g., Busch et al., 2009; van Dijk et al., 2008; Drewes & VanRullen, 2011; Ergenoglu etal., 2004; de Graaf et al., 2015; Gonzalez-Adino et al., 2005; Hanslmayr et al., 2007; Melcón et al., 2024; Romei et al., 2008; 2010; Thut et al., 2006; VanRullen et al., 2011; Zazio et al., 2021). In general, increased alpha power is especially – but not exclusively – related to poorer task performance (with a potential role of phase remaining controversial; Busch et al., 2009; Schroeder & Lakatos, 2009; VanRullen, 2016; Benwell et al., 2017; Bompas et al., 2015: Melcón et al., 2024; Ruzzoli et al., 2019; Zazio et al., 2021). Most studies use low-contrast stimuli evoking low detection rates or experimental conditions evoking exogenous variability, limiting comparisons to the current paradigm. We identified positive associations of endogenous behavioural variability with delta– and beta-band activity, generalising previous conclusions based on single-trial simple and choice RT (Bompas et al., 2015; Gonzalez-Andino et al., 2005), but did not find the previously reported correlations with alpha or gamma activity (Bompas et al., 2015; Paraskevopoulou et al., 2021). More importantly, prior studies reported that oscillatory activity explains only a small proportion of behavioral variance (∼1-8%), and our estimates fall at the lower end of that range.

To our knowledge, functional connectivity underlying within-subject variability have previously been examined only with fMRI (Kucyi et al., 2017; Rosenberg et al., 2020; Seeburger et al., 2024; Zuberer et al., 2021), despite calls for neurophysiological studies capable of tracking dynamic connectivity with higher temporal resolution (Kucyi, 2018). Our study therefore provides the first MEG evidence linking functional connectivity to within-subject variability. We note functional connectivity as derived by fMRI and MEG are not necessarily related, but amplitude-envelope-correlation measures as reported here have been shown to produce similar networks to fMRI-FC measures (Brookes et al., 2011). Indeed, our results are consistent with dynamic fMRI findings of increased within– and between-network connectivity as variability increases (Kucyi et al., 2017). Furthermore, we find that functional connectivity predicts behavioural variability independently of time-on-task. A similar independence of time-on-task was previously reported after detrending behaviour (Bompas et al., 2015), although our analysis controls for temporal trends in both neural signals and behaviour, providing a more stringent test of this relationship. The reported findings concern whole brain analysis, reflecting calls to move on from the canonical DMN network (Kucyi, 2018). Although network-specific analyses of DMN and task-relevant auditory-motor regions produced effects in the same direction, they explained less variance than whole-brain connectivity. Furthermore, connection-level analyses indicated that the observed associations were broadly distributed across the whole brain rather than driven by a particular subset of connections or networks.

We found that blink duration, pupil size variability, and head movement – but not mean pupil size – also predicted behavioural variability. Head movement is rarely examined except as a data quality criterion, although it has been used as a covariate in functional connectivity analyses (Rosenberg et al., 2020; Zuberer et al., 2021). In contrast, increased blink rate and duration (or ‘eyelid closure’) are common proxies for fatigue or sleepiness (Pan et al., 2025) or cognitive load related to off-taskness (Hollander & Huette, 2022). Behavioural correlates are more frequently reported in driving (e.g., Abe et al., 2011). Blinking itself strongly influences visuomotor RT when occurring around stimulus onset (Johns et al., 2009). Such effects are unlikely to generalise directly to continuous tasks without a clear visual component, such as the current paradigm. Nevertheless, our results suggest that associations between blink duration and behavioural variability extends beyond strict visuomotor domains.

Analyses of pupil size typically distinguish between ‘tonic’ (baseline/pre-stimulus) and ‘phasic’ (stimulus-evoked) activity. Our current findings relate to tonic activity, capturing ongoing fluctuations during event-free periods. Our findings mirror Martin et al., (2022), who reported an absence of within-subject relationships between tonic pupil size and RT in psychomotor vigilance tasks. In contrast, studies using binned or median-split RT have reported larger tonic pupil sizes for the slowest RTs both in the SART (Unsworth & Robinson, 2016) as well as auditory detection (Gilzenrat et al., 2010), associated with increased drift rate variability (Murphy et al., 2011). However, these studies relied on binned or median-split RT measures, which may account for the divergence from our continuous analyses. Time-on-task may be another key factor: van den Brink et al. (2016) found that controlling for time-on-task changed the predictive shape of pupil size for RT variability and slowest RT on a CPT from linear to quadratic. We found an attenuation of the within-subject linear correlations between behavioural variability and pupil size variability or cumulative blink duration, but they remained statistically present. As the oculomotor patterns remain after controlling for time-on-task for behaviour but not for subjective attention, they are unlikely to reflect time-dependent general task disengagement or fatigue.

#### Univariate predictors of subjective attention

We identified time-independent positive associations with theta-, alpha-, and beta-band power. Positive associations with alpha have been particularly reported in the literature (e.g., Baldwin et al., 2017; Compton et al., 2019; Jin et al., 2019; Kam et al., 2021; Macdonald et al., 2011; Polychroni et al., 2022; Whitmarch et al., 2014; 2017). Delta has also been highlighted as a positive predictor (Andrillon et al., 2021; Kam et al., 2022), but is reported infrequently. Although we found a positive association with delta power, it did not survive after correcting for time-on-task. A systematic review has suggested that power in the delta, theta and alpha bands are higher and beta power is lower during off-task states, and that the attentional locus on the task (i.e., internal orientation, such as during a breathing task, versus external orientation, such as during a visual task) may influence the directions of effect (Kam et al., 2022). However, directions of effect are inconsistent even within the same “types” of tasks, and many studies reported non-significant results. Recent evidence suggests that alpha has differing effects depending on whether people have their eyes open or closed (Thielking et al., 2026). It is also common for EEG research to selectively analyse and/or report one or two frequency bands to examine the relationship between neural activity and their dependent variable – with alpha being highly favoured. This prevalence may be explained by two constraints: 1) EEG’s reduced sensitivity at higher frequency, and 2) short time windows for within-trial analyses imposing a lower bound on frequency analyses. However, oscillatory power estimates across frequency bands are not statistically independent, as reflected in amplitude correlations between frequency bands and shared temporal dynamics (e..g, Carlqvist et al., 2005; Godfrey & Singh, 2021; Linkenkaer-Hansen et al., 2001). Overall, our results do not support a single-minded focus on alpha as an oscillatory marker of attention, as off-taskness was preceded by increases in multiple bands, and oscillatory power were highly correlated within subjects across frequency bands.

We further identified weak correlations between off-taskness and pupil size, pupil size variability, blink duration and head movement, although only the pupil size variability remained after controlling for time-on-task. Our results echo previous reports of positive associations between blink rates and off-taskness (Grandchamp et al., 2018; Smilek et al., 2010), also gone when controlling for time-on-task (Steindorf & Rummel, 2020). Results on tonic (baseline) pupil sizes have been highly inconsistent – reporting increases (e.g. Franklin, 2013; Jubera-Garcia et al., 2020; Groot et al., 2021; Unsworth et al., 2019), decreases (e.g. Gouraud et al., 2018a; 2018b; Grandchamp et al., 2018; Huijser et al., 2018; Unsworth & Robison, 2016; 2018), or no significant effects (Groot et al., 2022; Huijser et al., 2020; Jubera-Garcia et al., 2020). A recent review paper identified three reasons for inconsistencies: 1) heterogeneity of probe-based off-taskness, 2) large array of tasks differing in multiple dimensions including cognitive processes, locus of attention, and effort, 3) analysed time windows preceding off-taskness reports ranging from less than a second to 1-2 minutes (Pelagatti, 2024). Based on our findings, we would like to add a fourth: whether temporal trends are corrected for or not. We observed decreases in pupil size over time, consistent with previous reports (e.g., van den Brink et al., 2016; Huijser et al., 2020).

Our results so far treat off-taskness as a single construct without distinguishing ‘mind wandering’ and ‘mind blanking’. There is evidence suggesting that these are distinct states with different neural profiles (Andrillon et al., 2025; Mortaheb et al., 2022; Munoz-Muset et al., 2025; Stawarczyk et al., 2020) – which might account for the low predictive power of neurophysiological measures for subjective ratings observed in the current study. To explore whether our findings were sensitive to this distinction, we conducted additional analyses separating subjective off-task experiences into *mind wandering* and *mind blanking (drowsy)* on all participants who had at least two ratings in both categories (N=11) and found that preceding oscillatory states were highly similar. Although these analyses are strictly exploratory due to the low number of participants and the strong imbalance of trials between the categories, we conclude that it is unlikely that the low predictive power of neurological markers is driven by a mixture of neural profiles.

#### Multivariate prediction

Individual predictors accounted for small portions of variance. This is unlikely to reflect measurement imprecision or underpowered effects, but rather the expected multi-causal signature of moment-to-moment fluctuations. Univariate predictor-outcome pairs often may reach statistical significance, but unlikely provide a complete description. In line with this view, combining predictors across modalities yielded clear benefits for behavioural variability. Here we tested 13 different predictors from three categories. For behavioural variability, the three predictor-categories had an independent contribution – with the model combining all predictors clearly outperforming each single model, even after accounting for variance explained by chance, even though some shared variance was present.

These findings replicate and expand on previous literature. For example, pupil size and EEG-derived entropy have been shown to correlate within subjects with low shared variance (1%) in a pitch discrimination task (Waschke et al., 2019). Their relationships to RT were linear and quadratic in shape respectively – showing a benefit of combining predictors. Similarly, Podvalny et al. (2021) showed that associations between pupil size and MEG-based oscillatory power were widespread across networks and frequency bands. While binned RT was primarily associated with pupil size and visual delta power, broader associations emerged for signal detection measures such as sensitivity (d′). Model comparisons indicated that a joint model of oscillatory power and pupil size performed best, although their contributions were not unique. Notably, neither paper accounted for time-on-task, and both have elements of exogenous variance (varying stimulus levels and inter-trial intervals). Building on these results, our current findings show a multi-modal contribution on purely endogenous fluctuations that remains even after detrending.

For off-taskness, the benefit of each separate modality was less evident because effects of time-on-task overlapped with the effects of neurophysiology, to the extent that the correlations between overt physiology and off-taskness were mostly gone after detrending. Previous work has likewise shown that multiple physiological and temporal variables can jointly predict subjective off-taskness ratings, including pre-stimulus pupil size, alpha power, and experimental block (Whitmarsh et al., 2021).

However, mediation analyses suggested that apparent pupil effects were mediated by alpha power – but these mediation analyses did not consider temporal effects. Machine-learning approaches have also demonstrated improved classification of off-versus on-task states as additional modalities are included, spanning EEG-derived pre-stimulus oscillatory power, fMRI-derived connectivity, tonic plus phasic pupil size, and evoked potentials (Groot et al., 2021). However, connectivity was the dominant predictor, with limited clarity regarding unique contributions from other modalities. Again here, time-on-task was not considered. Together, our findings highlight that multimodal prediction is most informative when shared temporal structures are removed.

Complicating the wider picture, cross-predictor correlations may reflect similar underlying mechanisms (e.g., arousal; Podvalny et al., 2021; Waschke et al., 2019) but may also reflect artifacts. While interesting science happens outside the “modality islands” from Figure 4, the strongest correlations likely reflect such artifacts – such as the correlation between functional connectivity and head movement. This highlights the importance of testing whether each modality adds predictive value. While we cast a wide net in the current study, there are many other predictors to consider, including but not limited to heartbeat (e.g., Park et al., 2014; Salomon et al., 2016), respiration (e.g., Johannknecht & Kayser, 2022), and stomach activity (e.g., Loescher et al., 2026; Rebollo & Tallon-Baudry, 2022; Richter et al., 2017), as well as a plethora of other MEG measures. Still, we hope the current paper can provide a new perspective on combining multi-modal measures. This includes controlling for temporal effects as well as estimating explained variance as an effect size – which remains underreported in the field, despite strong evidence that the absence of effect-size reporting undermines interpretability, comparability, and cumulative science (e.g., Li et al., 2023; Szucs & Ioannidis, 2017) – while controlling for variance explained by chance (Perquin et al., 2024).

### Time-on-task is a key player

Establishing causality among multiple correlated predictors is challenging. Time-on-task, however, is an unusual case: unlike many neurophysiological or behavioural measures, it can only act as a cause and not as an effect. Its effect on both behavioural variability (e.g., Groot et al., 2022; Perquin et al., 2020; 2023; 2024; Torre & Wagenmakers, 2009; Wagenmakers et al., 2004; Zanesco et al., 2023) and subjective off-taskness (e.g., Groot et al., 2022; Irrmischer et al., 2018; Thomson et al., 2014; Zanesco et al., 2023; 2025) is well-established. Yet, explicitly controlling for it in statistical analyses remains uncommon. As highlighted throughout the current paper, controlling for temporal trends is important to appreciate both univariate predictor-outcome pairs as well as multi-modal analyses. Detrending does not necessarily attenuate correlations: the relationship between neurophysiological measures and behavioural have been reported to persist (Benwell et al., 2017; Bompas et al., 2015; Kopčanová et al., 2025), disappear (Bompas et al., 2015; Kopčanová et al., 2025), or change predictive form (van den Brink et al., 2016). In our results, it revealed a correlation between beta power and behavioural variability that was inconclusive before detrending.

Our results show temporal trends in pupil size and other oculomotor variability, in agreement with previous literature (van den Brink et al., 2016). We also found increases over time in head movement and functional connectivity, but not on the oscillatory MEG measures. Previous studies have found effects of time across theta, alpha, and beta (Benwell et al., 2019; Kopčanová et al., 2025; Stol et al., 2016). One explanation may be that linear temporal effects are small, so different analysis types may push them in or out of statistical significance. Indeed, Benwell et al. (2019) report a shared variance of about 1.5%. In practice, they are unlikely to be zero – especially not on the level of the individuals. Therefore, detrending is important even if a statistical threshold is not met, for improved characterisation of intra-individual dynamics.

### Individual differences

If one aims to predict people’s upcoming performance, the group average is only useful in so far as it reflects the patterns of the individuals. Our findings show that there is not a single marker nor a robust set that fits all even for those numerically strongest and most robust on the group level. Substantial inter-individual variation in predictors has been described previously (Dhindsa et al., 2019; Jin et al., 2019; Kucyi et al., 2025; Nakuci et al., 2023) and may have clinical implications (Tamm et al., 2025). Our and other results indicate that group averages can obscure individuals, and more importantly, that it is unlikely that we will find universal signatures for either subjective attentional state or behavioural variability.

### Conclusion

Behavioural variability and subjective attentional state have been considered as different markers of the same underlying processes. Yet, on the metronome task, their shared variance is low and their neurobiological predictors are dissociable. If there are any internal states that we have access to, these are not the states underlying moment-to-moment fluctuations in our behaviour. The idea that these are “two sides of the same coin” may thus reflect intuitions about subjective experience rather than the structure of underlying neurocognitive mechanisms.

## Methods

### Participants

Twenty-one participants (eighteen female, 3 male, 21-40 old, *M_age_* = 26.3) were tested in a two-session experiment. All of them had normal or corrected-to-normal vision and hearing, and no psychiatric or neurological disease. Participants were paid £10/hour (excluding potential reward, see below). The study has been approved by the Cardiff University School of Psychology ethics committee. As the current study is focused on within-subject fluctuations from trial to trial, statistical power here concerns the number of observations per participant. Sample size was constrained by time. Due to technical issues, one participant only completed five blocks out of six, and for one other participant, the behavioural responses were not recorded on three blocks out of six.

### Materials

The experiment was generated outside the magnetic room using MATLAB version 8 (The Mathworks, Inc., Release 2015b) and Psychtoolbox-3 (Brainard, 1997; Kleiner et al., 2007; Pelli, 1997), with a HP Z230 Workstation PC, and was displayed to the participants on a screen inside the magnetic room with use of a PROPixx DLP LED projector. The background was set at light-grey with a white fixation dot in the centre. Participants were seated 185 cm away from the screen, with their head on a chinrest to limit head movement. Auditory stimuli were delivered to earplugs via MEG-compatible tubing connected to an air-puff stimulator. The metronome sound consisted of a sharp transient click lasting 75 ms. Manual responses were recorded with a Nata Technologies 2-hand Button System. Head localisation was conducted before each block. Standard fiducials were applied to the participant’s left and right tragus and eye centre.

Whole-head MEG activity was recorded at 600Hz with the CTF-Omega 275 channel radial gradiometer system (VSM MedTech). Eye movements and pupil diameter were recorded binocularly at 500 Hz with an Eyelink 1000. Head location was also tracked continuously during the experiment, using the 3D coordinates of the three head fiducials.

### Procedure

Rhythmic reaction time (RT) was measured on each trial on an adjusted version of the Metronome Response Task (MRT; Seli et al., 2013). Each trial lasted 3000 ms, and 1500 ms after trial onset, a short tone of 75 ms was presented. Participants were instructed to press in synchrony with the tone – aiming for a response time of 0 ms in relation to the tone on each trial.

Subjective ratings of performance and off-taskness were both measured with quasi-randomly presented probes on a scale from 1 to 9 throughout the task (Figure 6). Each probe consisted of four consecutive subprobes. The first two subprobes referred to their subjective off-taskness (*“Please record a response from 1 to 9 which characterises how on task you were just before this screen appeared”*, with 1 as ‘completely ON task’ and 9 as ‘completely OFF task’) and performance (*“Please record a response from 1 to 9 which characterises how you rate your performance before this screen appeared”*, with 1 as ‘excellent’ and 9 as ‘terrible’). The order of these two probes was counterbalanced over sessions and participants. With the current phrasing, we expect the relationships between SDRT, off-taskness, and subjective performance to always be positive, with higher values indicating worse states.

**Figure 6.**
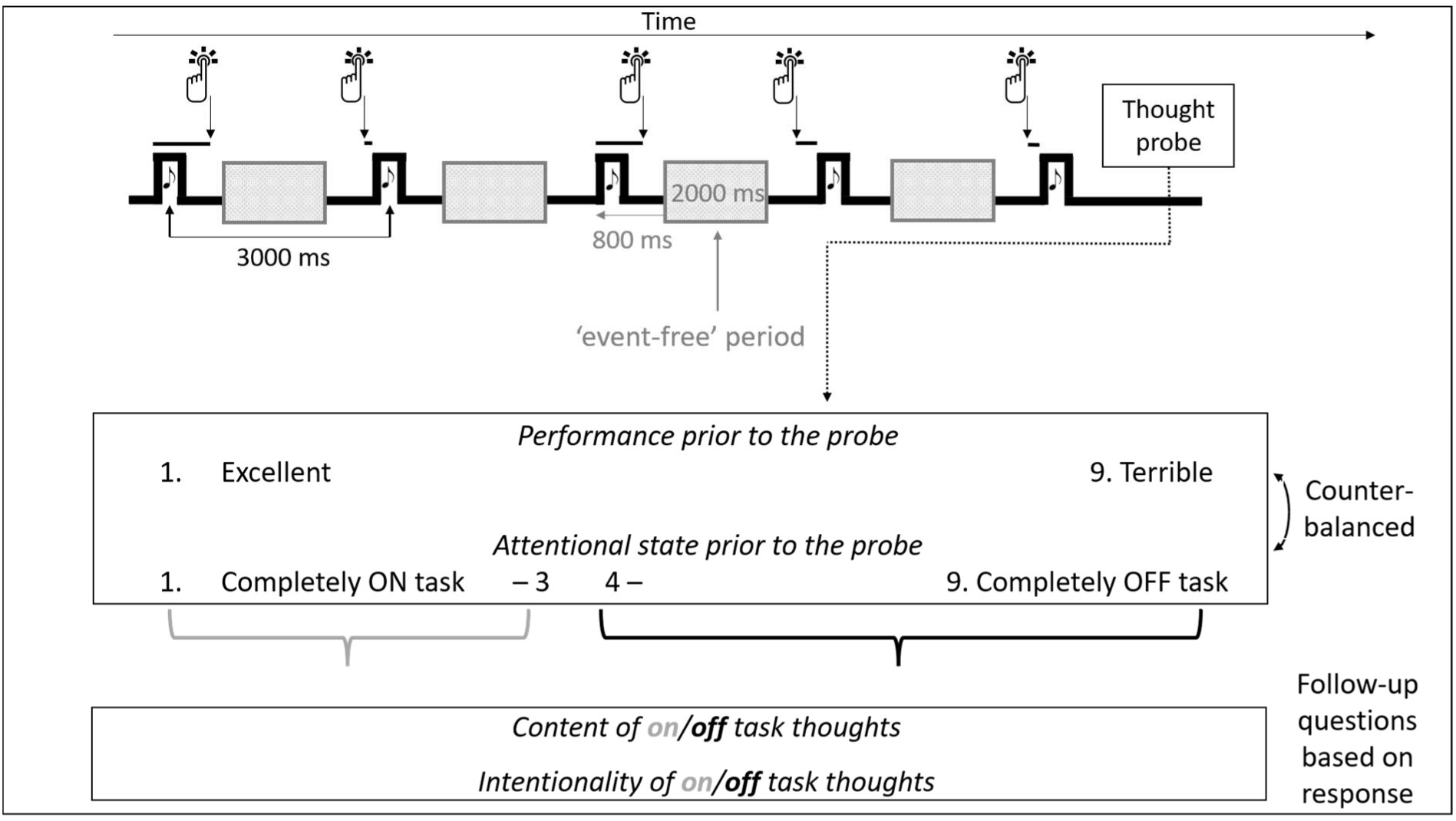
Overview of the Metronome task, and the four probes with which participants were presented quasi-randomly throughout the task. Participants were instructed to press in synchrony with a tone, occurring every three seconds, and their relative reaction times were recorded. The 2-sec ‘event-free periods’ (starting 800 after tone-onset) target the periods between the tones and presses, in order to estimate ongoing physiological and neural activity.

Next, the participants were presented with two follow-up questions on content and intentionality, depending on their off-taskness rating. Any rating above 4 was considered ‘off task’, and was followed by the same question on content with different response options – 1 as ‘bodily states’ (e.g., thinking about being uncomfortable in the MEG), 2 as ‘mind wandering’ (i.e., engaged in task-unrelated thoughts), 3 as ‘mind blanking – alert’ (i.e., thinking about nothing, but still in an alert state), and 4 as ‘mind blanking – drowsy (i.e., thinking about nothing due to being too tired/falling asleep) – and a question on intentionality (*“Would you say your off-task thoughts were: 1. Intentional: Conscious off-task thoughts, or 2. Unintentional: Off-task thoughts despite your best efforts”*). Any rating between 1 and 3 was considered ‘on task’, and was followed by the questions: (*“Please indicate with the corresponding number what you were thinking about specifically?”*, with 1 as ‘the tones’, 2 as ‘the key presses’, 3 as ‘the instructions of the task’, and 4 as ‘your performance on the task’) and (*“Would you say your on-task thoughts were: 1. Intentional: Conscious on-task thoughts, or 2. Accidental: On-task thoughts without your best efforts”*). These were added with no other purpose than balance the number of questions following low and high off-taskness reports. Note that the “on-task” and “off-task” categories were only relevant to determine content and intentionality questions, while on-taskness and off-taskness throughout the manuscript refer to a continuum from 1 to 9.

The experiment consisted of two sessions, both lasting about 1.5 hour, with the same structure: each consisted of 3 blocks of 240 trials, resulting in (2 * 720 =) 1440 trials per participant. Each of these blocks contained 8 thought probes, resulting in (2 * 24 =) 48 thought probes per participant. Each probe was presented on a random trial within each 30-trial sub-block, excluding the first and last five trials of each sub-block – to ensure the probes were not administered too close together (i.e., less than 10 trials apart). These are critical choices in the current design: the sub-blocks cannot be too short, as too many interruptions would disrupt our measure of RT variability, but also not too long, as that would result in underpowered analyses for the subjective ratings. Prior to the main experiment, participants conducted a training of 60 trials, with a thought probe being presented after trial fifteen. After the training, the experimenter checked the performance to ensure the participant understood the task and gave the participant feedback on their performance.

To give the participants motivation to perform as well as possible throughout the task, we used the random lottery reward system (Cubitt et al., 1998). At the end of a session, one random trial number was extracted. If the moving window standard deviation on this trial was lower than .075, the participant received £5 extra. This cut-off was based on pilot data, such that the chance of reward would be ∼20%.

Before and after the task, participants conducted a 4-minute resting state, in which they were instructed to fixate on the dot in the centre of the screen. Afterwards, participants completed questionnaires on daydreaming (Daydreaming Frequency Scale; Singer & Antrobus, 1963), ADHD tendencies (Adult ADHD Self-Report Scale v1.1; Kessler et al., 2005), and impulsivity (UPPS-P Impulsive Behaviour Scale; (Whiteside & Lynam, 2001; Lynam et al., 2006). The eye tracking data during rest alongside the questionnaire scores have been published previously in Perquin & Bompas (2019).

### Data preprocessing and analysis

#### Task-related measures (non-physiological)

*Reaction Time Variability.* Reaction time variability (SDRT) was calculated on every trial *n* as the standard deviation across the RT from trial *n-4* to *n* as per Seli et al. (2013). This results in a moving window measure of variability from trial to trial. With each trial lasting 3 seconds, this resulted in time window of 15 seconds. The first five trials of the session as well as the first five trials after each thought probe were excluded from analysis. Contrary to previous work (Anderson, Seli, Perquin), the SDRT was not logarithmically transformed within participants, because the distributions were closer to normal for the untransformed data. This discrepancy might be due to the longer time delay between tones.

*Previous RT variability.* To estimate short-term temporal dependency in the RT series, previous SDRT was calculated on every trial *n* as the standard deviation across the RT from trial *n-9* to *n-5*. As such, SDRT and previous SDRT had no overlap in responses – mimicking an autocorrelation.

*Time-on-task.* Time-on-task was operationalised as blockwise trial number, ranging 1-240 for each of the six blocks (Bompas et al., 2015, Perquin et al., 2024), ignoring interruptions caused by thought probes. This allowed for examination of longer-term temporal linear trends.

*Subjective ratings of off-taskness and performance.* Before conducting any analyses, we were interested in the frequency with which participants reported being on-versus off-task, as well as the content of their off-task thoughts. Figure 7 shows the percentage of on-task reports over the two sessions, as well as the breakdown into percentages of off-task reports, at the subject and group levels. Particularly striking is the large variability within and between participants. When they reported being off-task, ‘mind wandering’ was the most common category. ‘Drowsy mind blanking’ and distractions due to ‘bodily sensations’ were rarer, but did occur. One participant never reported being on-task, and another participant only reported to be off-task once throughout both sessions. These two participants were excluded from all analyses that relied on fluctuations in off-taskness ratings (dotted lines in Figure 7).

**Figure 7.**
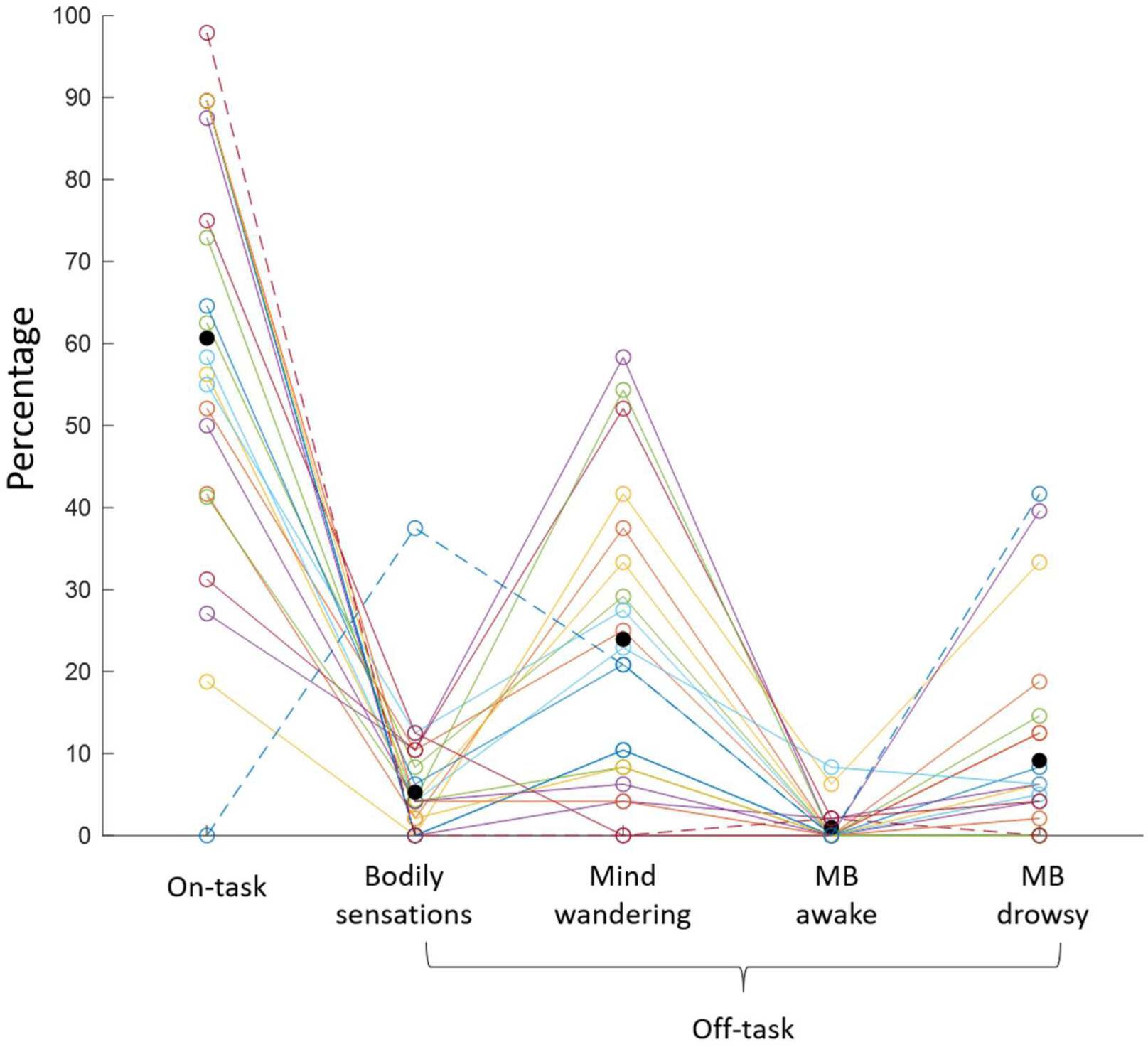
Percentages of on-versus off-task thoughts pooled across the two sessions, with off-task thoughts broken down according to content as reported in the follow-up question. Shown are both the individual participants (with each coloured line showing one participant. Two participants were excluded from all analyses involving subjective ratings (represented by the dotted lines). The black dots show the group medians in each category after exclusion.

It is clear that not all off-taskness is mind wandering per se – the other categories are less prominent but still a substantial proportion altogether. There was not enough variance in participants’ answers to analyse the different types of off-taskness separately. Therefore, we used all ratings regardless of content as a measure of subjective off-taskness. Likewise, intentionality could not be analysed as reports nearly all fell within two categories: intentional on-taskness or unintentional off-taskness. Such patterns have been found previously (Perquin et al., 2023; Unsworth & Robison, 2019), though others have reported intentional off-taskness (e.g., Golchert et al., 2017; Kane et al., 2021). We associated both subjective performance and off-taskness ratings with the five trials immediately preceding each thought probe. For example, if a thought probe appeared after trial n and the participants rated their off-taskness as ‘7’, we assumed that their state was ‘7’ throughout trial n-4 up until n.

For exploratory control analyses on content (see Discussion), we selected participants who reported both mind wandering and mind blanking (drowsy) at least twice (N = 11). Analyses were run separately for ‘mind wandering’ (on-task ratings [1– 3] plus off-task ratings [4–9] classified as mind wandering) and ‘mind blanking’ (on-task ratings plus off-task ratings classified as ‘mind blanking drowsy’).

*Event-free periods*. We extracted ‘event-free’ periods of two seconds for each trial, to investigate physiological and neural activity independent from activity related to the tones or button presses (see Figure 6). Because participants were inclined to respond before the tone, these event-free periods ranged from 2800-800 ms before each tone. All measures specified below have been estimated within this window.

To match the time scale of the physiological and neural measures to that of SDRT and subjective ratings, we calculated the mean across trials *n-4* to *n* for every trial *n* for each measure.

#### Overt physiological measures

*Cumulative blink duration.* Blinks were defined as missing eye-tracking data. Because the distribution of blink rates was narrow and skewed, we choose to analyse the cumulative blink duration. Therefore, no maximum duration was set for blink extraction. The duration of each blink during the event-free period from trial n-4 to trial n were summed to obtain a measure of cumulative blink duration for each trial n. This measure thus reflects how long participants had their eyes closed over the event-free period from the last five trials. Note that for three participants, the eye tracking data of the second session were not recorded due to technical issues.

*Pupil size mean and variability.* To minimise noise, any samples occurring 20 ms before or after a blink were excluded prior to the analysis of the pupil data. To correct for differences in pupil size across the two sessions, we first corrected all the samples during the task with the mean pupil size during the four-minute resting state preceding the task (see Perquin & Bompas, 2019) separately for both sessions. Because the relationship between pupil size and behaviour is complex (see Discussion), we quantified both the mean pupil size as well as the variability (standard deviation) in pupil size.

*Head movement.* To obtain one measure of head movement within a trial, the standard deviation of the 3D coordinates for each of the three fiducials was calculated for each trial, and then averaged across the fiducials.

#### MEG measures

Prior to beamforming, all MEG sensor data had a 1 Hz high-pass and a 150 Hz low-pass filter applied. The data was then epoched into trials and visually inspected so that trials containing large artefacts relating to eye and/or head movements could be identified and removed from the subsequent analyses. MEG activity was co-registered to structural MRI scans using the fiducials, which was verified with photographs. The analysis pipeline for denoising, co-registration, beamforming, reduction to the 90-node AAL atlas largely followed Koelewijn et al. (2019), adjusted to provide trial-by-trial estimates.

*Oscillatory power.* Localisation of MEG sensor data was determined with a linearly constrained minimum variance (LCMV) beamformer (6mm grid, single-shell forward model, Nolte, 2003) using FieldTrip version 20161011 (Oostenveld et al., 2011) to determine the beamformer weights for each voxel (in brain space from the structural MRI) in six frequency bands (delta: 1-4, theta: 3-8, alpha: 8-13, beta: 13-30, low gamma: 40-60, and high gamma: 60-140Hz). Subsequently, beamformer weights for each band were normalized (as in Hillebrand et al., 2012). This was done separately for different scanner sessions acquired for the same participants so that head localisation was consistent.

The source-space data from the different sessions were concatenated and reduced to 90 nodes (from the AAL atlas, Tzourio-Mazoyer et al., 2002) by selecting the virtual sensor in the AAL region with the greatest temporal standard deviation. The reduced 90-node virtual time series were segmented such that only the event-free epochs were included. This gave a time series for each of the 90 nodes within the AAL atlas separately for each trial. Finally, the mean oscillatory power was calculated for each trial separately for each frequency band.

*Functional connectivity.* Amplitude envelopes for each node time course were evaluated using the Hilbert transform. Symmetric orthogonalization was applied to the time series to avoid spurious correlations due to leakage (Colclough et al., 2015). One connectivity matrix was determined for each participant at each trial, meaning that changes in connectivity could be correlated with behavioural measures within participants. Functional connectivity between regions for each frequency band was defined as the Pearson correlation coefficient between amplitude envelopes across time within a single trial. A Fisher transform was used to obtain the z-scores from the Pearson correlation coefficients (Koelewijn et al., 2019).

This provided a z-scored 90|90 connectivity matrix for each trial and each frequency, separately for each participant. A combined measure of functional connectivity was then obtained by taking the vector sum of the connectivity matrix within each frequency band.

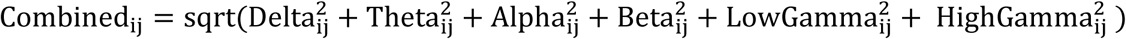

The advantage of this approach is that it increases the robustness of connections and increases the power of the analyses by reducing the number of comparisons (Koelewijn et al., 2019). Note that the connectivity is undirected and has no time-lag, so connectivity matrices are symmetrical (entry i,j is identical to entry j,i), this gives 4005 (90*(90-1)/2) unique connections within the 90×90 connectivity matrix. The mean connectivity strength was calculated for each trial by taking the mean of all connections between all nodes. We also explored whether specific frequency bands or networks were driving our results. Two networks were considered: the default mode network, hypothesised to be correlated with SDRT and off-taskness, and a task related network comprising the auditory and motor cortices, hypothesised to show the opposite pattern. For each network, nodes associated with this network were selected, and functional connectivity of all the pairs of nodes belonging to this network were averaged. The default-mode network comprised the posterior cingulate gyrus, superior frontal gyrus (medial), angular gyrus, and middle temporal gyrus. The task-related network comprised the precentral gyrus, supplementary motor areas, Heschl gyrus, and superior temporal gyrus.

#### Data analysis

Relationships between pairs of variables were computed using Bayesian Pearson correlation analyses (Figures 1-4). Distributions were compared to zero using two-sided Bayesian one-sample t-tests, with exception of the test to compare the overlap between predictors (histogram on Figure 2) – which was one-sided to test for the directional hypothesis. Note that for the overlap analyses (Figure 2), any negative coefficients were multiplied by –1, because we were interested in the strength rather than the direction of the correlation. Therefore, only positive correlations would be theoretically meaningful.

To test if temporal trends had any confounding effects on the relationship between other variable pairs, both variables were first regressed on blockwise time-on-task. We extracted the residuals from these regressions, representing variance independent of time-on-task, and subsequently recalculated the correlation across the residuals as above (Figure 4).

Multiple Regression analyses were conducted to extract the explained variance (R^2^) by combining multiple objective predictors. For both outcome measures, we ran four different models with: 1) task-related (or non-physiological), 2) overt physiological, 3) MEG, and 4) all predictors (Table 1). This provides us with a ‘ceiling of predictability’ of behavioural variance in the current task. The subjective ratings were not included as predictors for SDRT as they are measured ‘post-hoc’. Because estimates of explained variance are typically inflated due to overfitting (Yarkoni & Westfall, 2017), we corrected them for variance explained by noise (Perquin et al., 2024). First, we shuffled the outcome variable 100 times and ran the same regression model for each iteration – to obtain 100 estimates of R^2^_noise_. The corrected estimate (R^2^_corrected_) was calculated by subtracting the mean of these 100 estimates from the original R^2^. The explained variances reported in Figure 6 correspond to R^2^_corrected_.

**Table 1.** Overview of the predictors across the different regression models. *\*SDRT was included as a predictor in the models with off-taskness as outcome measure, as it is an objective marker measured preceding the thought probe*.

|  | Task-related | Physiology-model | MEG-model | Full model |
| --- | --- | --- | --- | --- |
| SDRT | * |  |  | * |
| Previous SDRT | X |  |  | X |
| Time-on-task | X |  |  | X |
| Pupil size |  | X |  | X |
| Pupil variability |  | X |  | X |
| Blink duration |  | X |  | X |
| Head movement |  | X |  | X |
| Delta power |  |  | X | X |
| Theta power |  |  | X | X |
| Alpha power |  |  | X | X |
| Beta power |  |  | X | X |
| Low-gamma power |  |  | X | X |
| High-gamma power |  |  | X | X |
| Function connectivity |  |  | X | X |

#### Bayesian statistics

All Bayesian statistics throughout the current research were calculated in JASP (JASP Team, 2017) using equal prior probabilities for each model and 10000 Monte Carlo simulation iterations. For BF_10_ > 1, higher BF_10_ indicate more evidence for the alternative hypothesis (i.e., the distribution is different from zero), and for BF_10_ < 1, lower BF_10_ indicate more evidence for the null hypothesis (i.e., the distribution is not different from zero). Across figures, we consider any BF_10_ between 1/3 to 3 to be indeterminate, consistent with common guidelines. BF_10_ > 10 and > 100 are considered strong and very strong. We do not distinguish ‘extreme’ evidence to reduce the visual information on the figures. The exact Bayes Factors can be found in the annotated JASP files.

### Data and code availability

The raw behavioural and eye tracking data, full analysis code, processed summary scores, and annotated JASP files will be made available on OSF after publication (reviewer link: https://shorturl.at/DUpvH). The raw and source localised MEG data will be made available after publication

### Author note

We would like to thank Courtney Taylor and Lingsi Zhou with their help in collecting the data.

MNP: conceptualisation, methodology, analysis, investigation, data curation, writing (original draft), visualisation, project administration.

MD: analysis, data curation, validation, writing (review & editing).

GP: supervision, analysis, resources, writing (review & editing).

KS: methodology, analysis, writing (review & editing).

AB: conceptualisation, methodology, analysis, writing (original draft), visualisation, resources, supervision, project administration.

## Supplementary Materials

To determine whether the widespread increase in connectivity associated with RT variability was driven by a subset of connections or was part of a nonspecific increase in connectivity, we correlated the strength of each connection separately. This gave a 90×90 behaviour-connection correlation matrix for each participant, where each entry (i,j) is the Pearson correlation coefficient between the strength of the connection (which connects AAL regions i and j) with RT variability. For example, entry (29,37) in the behaviour-connection correlation matrix for participant 1 would give the observed correlation between IIV and the strength of the connection between the left insula and the left hippocampus for that participant. T-statistics were determined for each behaviour-connection correlation by performing separate single-sample t-tests on each behaviour-connection-correlation. This involved 4005 t-tests (one for each unique behaviour-connection-correlation), where n=21 (number of participants). We ran the same analyses with the attentional state ratings.

To determine whether the t-statistics associated with each behaviour-connection correlation were significant, we performed a permutation test. The behavioural values for each participant were shuffled 100 times, and correlated with each connection, giving 100 pseudo-r values for each behaviour-connection correlation and each participant. These were then used to generate 100 pseudo-t-statistics for each behaviour-connection correlation. If the observed behaviour-connection-t-statistic was greater than the 95th highest pseudo-t-statistic this showed that the behaviour-connection correlation was significantly positive (α=0.05).

Figure A1 show histograms of the observed behaviour-connection correlation t-statistics (left: RT variability, right: attentional state ratings). Each dot is a behaviour-connection correlation where the significant values are red. The majority of both behaviour-connection correlations were positive, and a large proportion of these positive correlations were significant (RT variability: 93.6%, 42.7% significant; attentional state ratings:74.9% positive, 15.1% significant). The centres of the distributions for both measures were positive (i.e., there was an increase in mean connectivity). For both distributions, most connections were positive, and the distributions were approximately symmetrical, suggesting that no particular subset of nodes is driving the increase in connectivity. If the increase in connectivity was driven by a particular subset of nodes, then we might expect a distribution with two peaks, one at zero (those connections not related to behaviour), and a second smaller positive peak (those connections related to connectivity).

On the group level, we found that there was no clear pattern in the significant behaviour-connection (Figure A2; left) or attention-connection (right) correlations, nor any overlap to overlap with a particular canonical network or auditory-motor function. This along with the distributional plots suggests that the increase in mean connectivity is not driven by a specific subset. We found little overlap in significant nodes between RT variability and attentional state ratings.

**Figure A1.**
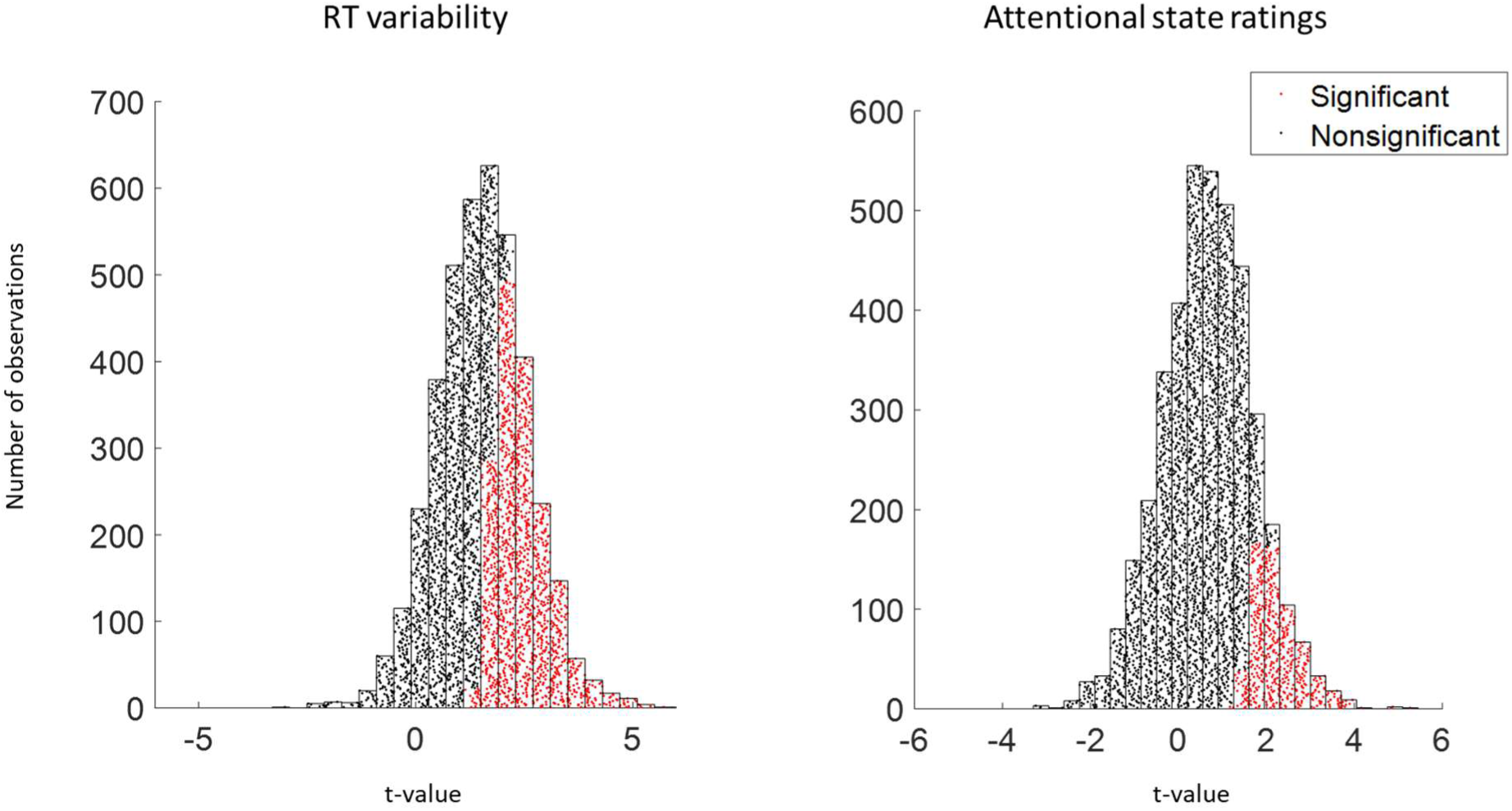
Histogram of the t-values associated with correlations between connection and RT variability (left) and connection and attentional state ratings (right). Black and red dots show those correlations which are nonsignificant and significant respectively on the group level.

**Supplementary Figure A2.**
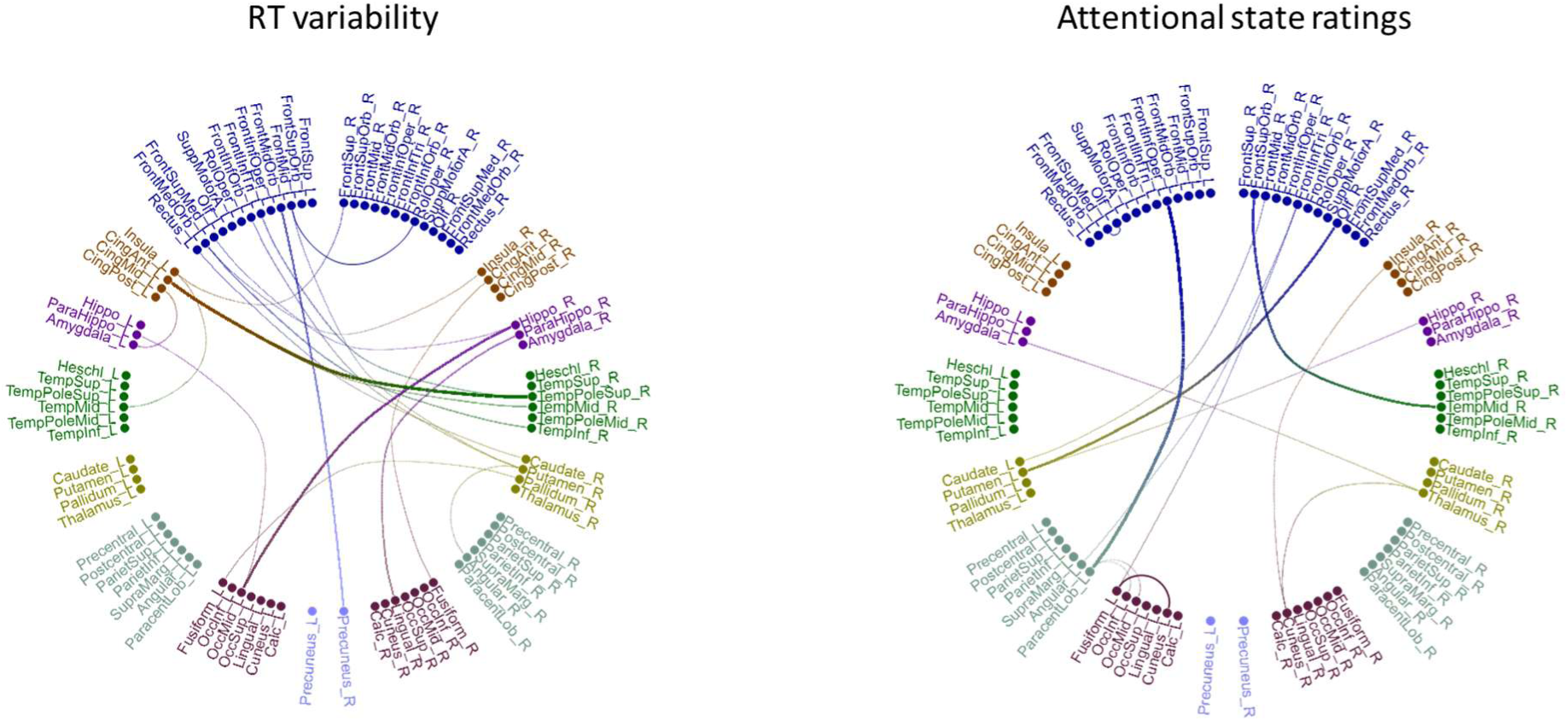
Web plot showing the significant behaviour-connection correlations, for RT variability (left) and attentional state ratings (right). Width of lines between AAL regions is determined by magnitude of t-statistic. There are no clear patterns, associated with canonical networks (e.g., DMN), and there is little to no overlap between RT variability and attentional state ratings.

## Footnotes

1 title and abstract contain: (performance OR “reaction time”) AND (metacognition OR “attention focus” OR “mind wandering”)

